# Accurate estimation of microbial diversity with Distanced

**DOI:** 10.1101/423186

**Authors:** Timothy J. Hackmann

**Affiliations:** Department of Animal Sciences, University of Florida, Gainesville, Florida, 32611, USA

## Abstract

Microbes are the most diverse organisms on Earth. Sequencing their DNA suggests thousands of different microbes could be present in a single sample. Errors in sequencing, however, make it challenging to estimate exactly how diverse microbes are. Here we developed a tool that estimates diversity accurately, even in the presence of sequencing errors. We first evaluated two existing tools, DADA2 and Deblur, which work by correcting sequencing errors. We found that these tools estimated within-sample (alpha) diversity poorly. In fact, we obtained better estimates if did not use the tools at all (left errors uncorrected). These tools performed poorly because they changed the relative abundance of different sequences; this is a side effect of correcting errors and discarding up to 90% of sequence reads in the process. Previous evaluations ignored sequence abundance when calculating diversity, overlooking this problem. Our tool, Distanced, differs from existing tools because it does not correct sequencing errors. Instead, it corrects sequence distances, which are used to calculate diversity. It does this correction with Phred quality scores and Bayes theorem. No sequence reads are discarded in the process. In our evaluation, Distanced accurately estimated diversity of bacterial DNA, fungal DNA, and even antibody mRNA. Given its accuracy, Distanced will help investigators answer important questions about microbial diversity. For example, it could answer how important is diversity for the planets ecosystems and human health.

## Introduction

Microbes are found nearly everywhere, and they form communities more diverse than any other group of organisms. Most communities likely host hundreds or thousands of different bacteria (ribosomal DNA [rDNA] sequences) [1]. Across all communities on Earth, there may be 10^12^ different species of microbes [2]. This diversity is not only fascinating, but it has important consequences for human health and the planets ecosystems. In the human gut, low diversity of bacteria has been associated with obesity [3]. In soil, low diversity of microbes has been associated with low plant productivity, nutrient cycling, and other measures of ecosystem function [4, 5].

Though microbial communities are no doubt diverse, it has been challenging to estimate exactly how diverse they are. Initial reports of a rare biosphere in seawater claimed unprecedented levels of alpha (within-sample) diversity. Most samples were estimated to have over 10,000 different bacteria (operational taxonomic units) [6]. Later analysis showed sequencing errors created false sequences, and the actual diversity was likely much lower [7].

Sequencing errors pose a problem for estimating diversity, but bioinformatics tools have been developed to tackle this problem. These tools aim to correct sequencing errors and output the original (error-free) sequences. DADA2 [8] and Deblur [9] belong to the latest generation of these tools, which claim accuracy to single nucleotide letters. Evaluations with artificial microbial communities would seem to support their accuracy and use for estimating diversity. In these evaluations, the number of sequences outputted by the tools closely matched the number of sequences (or organisms) known in the community [8–10]. When expressing diversity as the number of sequences, DADA2 and Deblur appear to have solved the problem posed by sequencing error.

Though DADA2 and Deblur estimate it accurately, the number of sequences (richness) is a very simple measure of diversity. Richness ignores how abundant or related sequences are, though these are important aspects of diversity [11]. If they use richness, investigators may miss ecologically important relationships in their data. For example, investigators found a relationship (positive correlation) between bacterial diversity and pH in a hot spring [12]. The relationship was weak when using richness as a measure of diversity. The relationship was strong, however, when using a more complex measure, mean pairwise distance. This more complex measure accounts for both sequence abundance and relatedness. Thus, evaluations of DADA2 and Deblur should consider not only richness, but also more complex ways of measuring diversity.

Our objective is to determine if bioinformatics tools accurately estimate diversity when accounting for abundance and relatedness. We find that DADA2 and Deblur do not. Indeed, they produced estimates worse than when the tools were not used (errors were left uncorrected). These tools corrected or removed most erroneous sequences, but they distorted sequence abundance in the process. We propose a tool, Distanced, that does not remove erroneous sequences. Instead, it corrects alpha diversity for the expected increase after sequencing, doing so directly with Bayes theorem.

## Results

We evaluated Distanced, DADA2, and Deblur using mean pairwise distance (MPD) as a measure of alpha diversity [13, 14]. As mentioned, it differs from richness by accounting for both abundance and relatedness (see also Fig. 1). It is calculated by averaging the distance between all pairs of sequences in a sample. Distance is defined here as the fraction of different nucleotide letters, but it can also be defined as the total number of different letters. Mean pairwise distance is also known as *θ*, and it is 1/2 the Rao Diversity Coefficient [11].

**FIG. 1:**
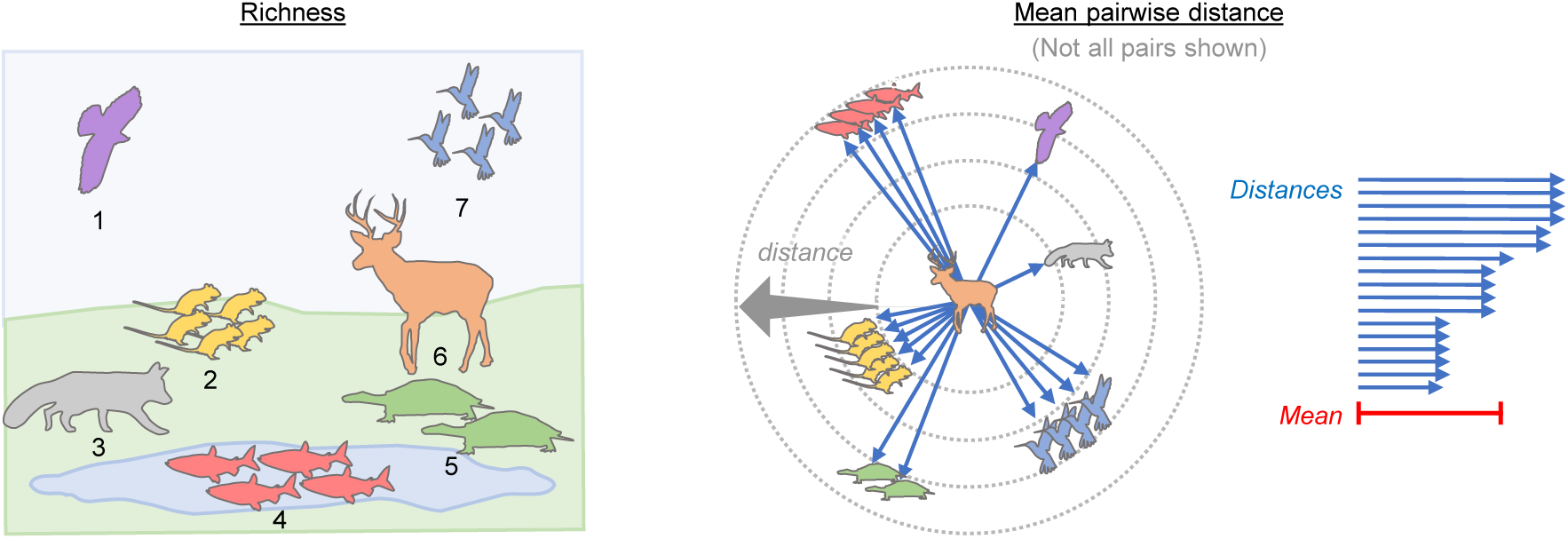
Comparison of richness and mean pairwise distance, which are two measures of alpha diversity. A non-microbial community is used for illustration. Distances are arbitrary.

Our tool estimates MPD before introduction of sequencing errors (Fig. 2). Sequencing errors inflate distances between sequences by changing their letters and making them more different. Our tool uses Bayess theorem to correct distances for introduction of errors (eq. [1] of Materials and Methods). The average of those corrected values is the estimated MPD. The only inputs required by the tool are 1) the observed distances (after introducing sequencing errors) and 2) error rates.

**FIG. 2:**
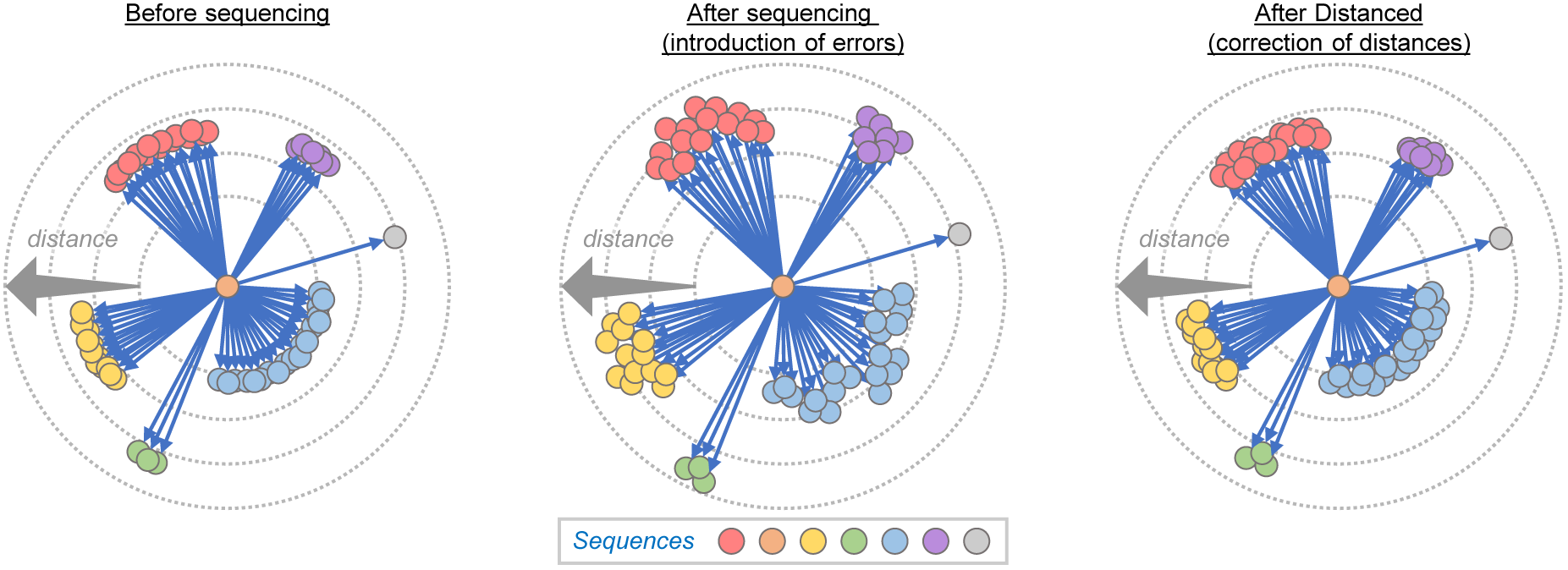
Approach used by Distanced to estimate alpha diversity (mean pairwise distance).

Error rates are the probability that a letter is incorrect after sequencing. If these rates are known, Distanced corrects distances exactly (without bias) and estimates MPD without error. This is demonstrated with simulated sequence reads (SI Appendix, Fig. S1).

In practical cases, error rates are not known, but they can be estimated by Phred quality scores reported by sequencing instruments [15]. Using these quality scores, we evaluated Distanced using rDNA sequences for bacteria and fungi.

We found that Distanced produces estimates of MPD close to their actual values, but estimates from DADA2 were generally worse (Fig. 3). Indeed, using no correction for sequencing errors generally produced better estimates than DADA2.

**FIG. 3:**
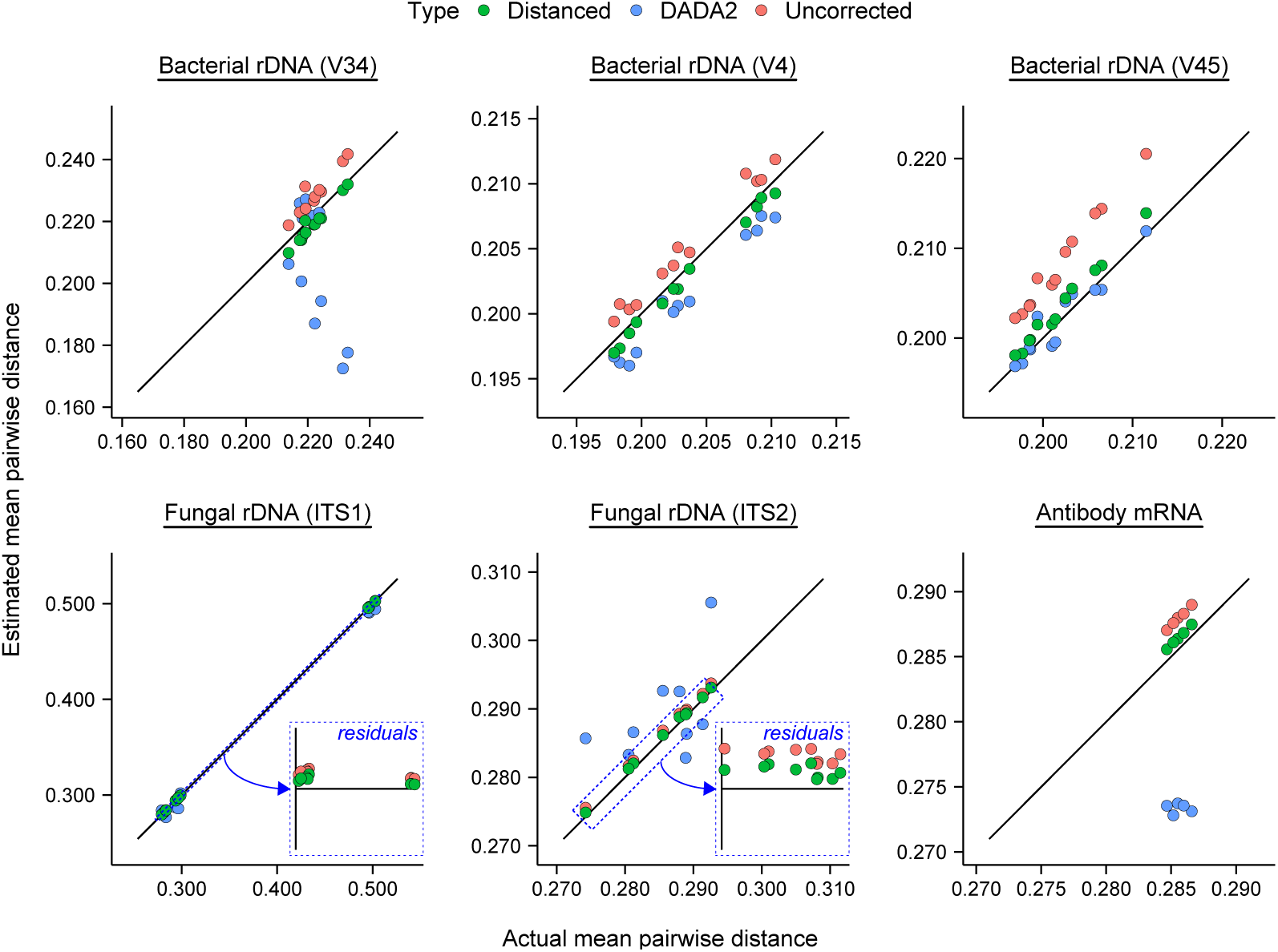
Performance of Distanced vs. DADA2 in estimating alpha diversity (mean pairwise distance) of ribosomal DNA (rDNA) from artificial microbial communities. Different regions (V4, V34, V45, ITS1, ITS2) are shown. mRNA from antibodies is included for comparison. Distances between sequences were corrected by Distanced, and errors in sequence letters were corrected by DADA2. Estimates of diversity when using no correction are shown for comparison. Full-length sequences were those analyzed. Each observation represents one sample.

Similar results were found when comparing Distanced and Deblur (Fig. 4). This comparison was made separate from the previous one because Deblur requires sequence reads that are truncated (trimmed to a fixed length).

**FIG. 4:**
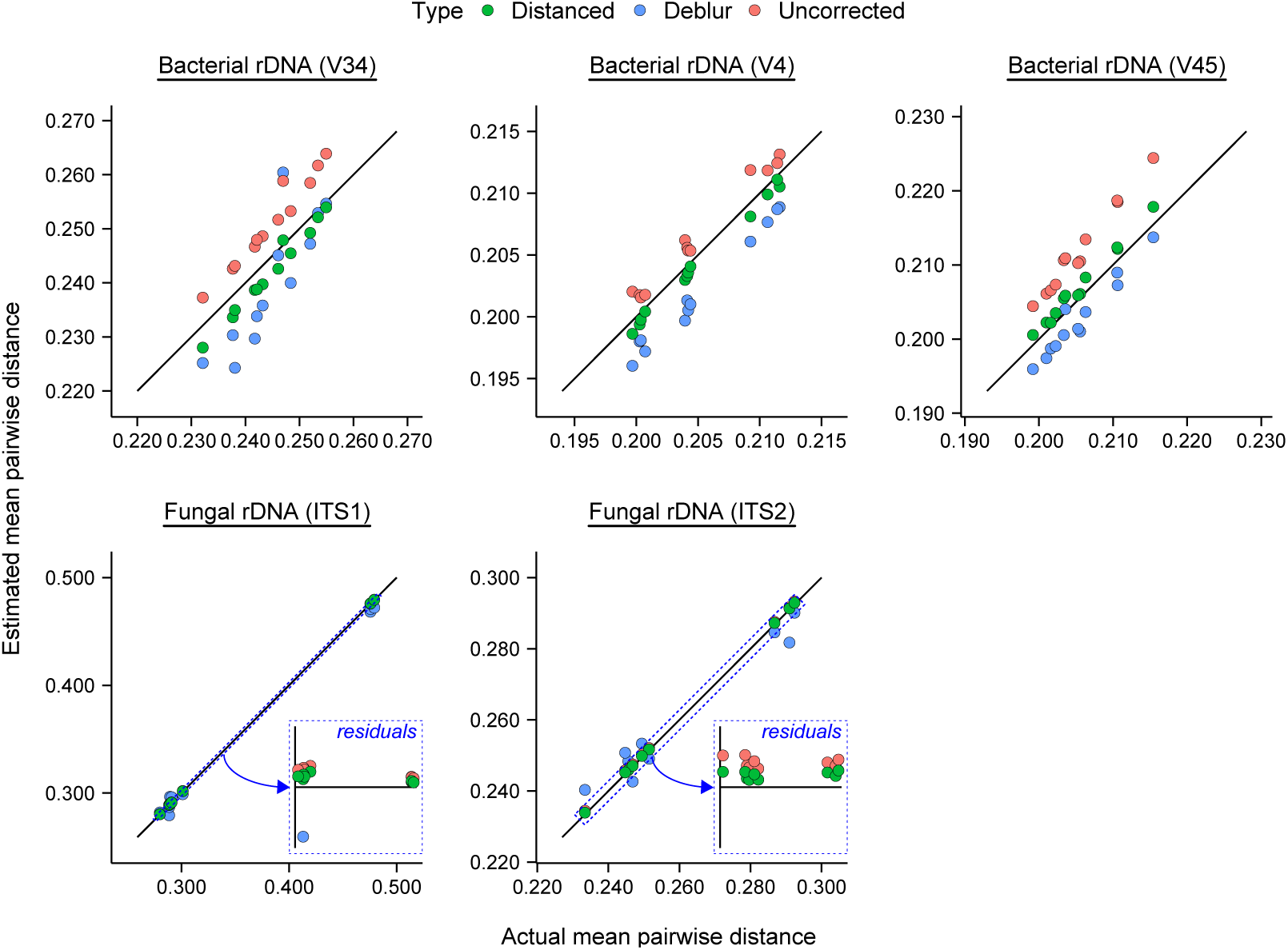
Performance of Distanced vs. Deblur in estimating alpha diversity (mean pairwise distance) of ribosomal DNA (rDNA) from artificial microbial communities. Errors in sequence letters were corrected by Deblur. Sequences truncated to a fixed length were those analyzed. See caption of Fig. 3 for further details.

We quantified performance of these tools by calculating root mean square prediction error. We found that Distanced always reduced error in MPD (Fig. 5). DADA2 and Deblur, in contrast, usually increased it.

**FIG. 5:**
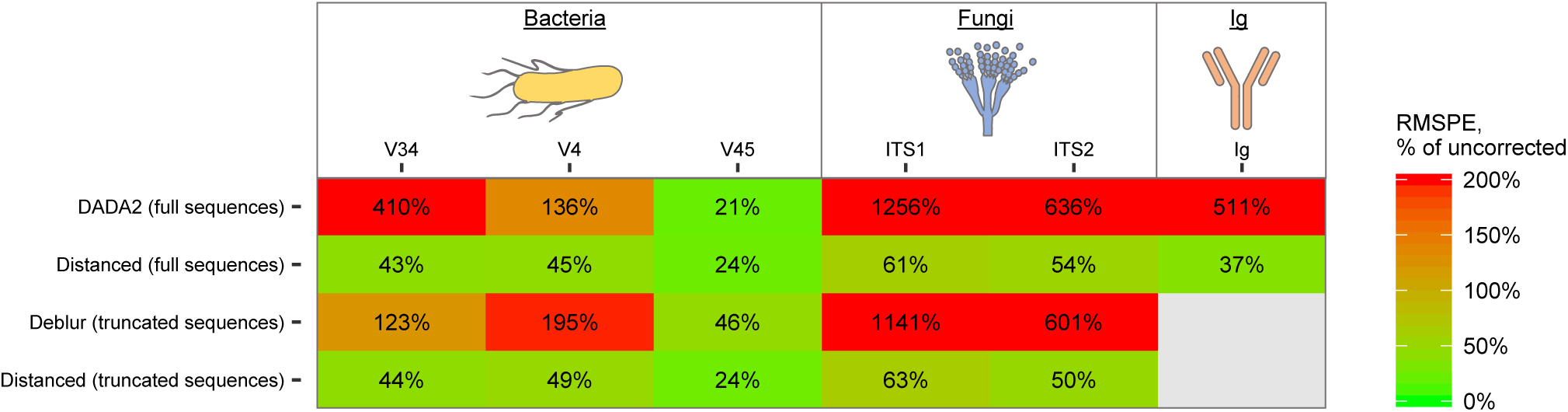
Error in estimating alpha diversity (mean pairwise distance) of artificial microbial communities and antibody mixtures by Distanced, DADA2, and Deblur. Root mean square error was calculated from observations in Fig. 3 and 4 and expressed as a percentage of using no correction for sequencing errors. A value > 100% means that leaving errors uncorrected is better. Ig = immunoglobulin (antibodies).

To determine why DADA2 and Deblur performed poorly, we first determined how many errors remained in the sequences they outputted. We found that almost no errors remained (Fig. 6 and SI Appendix, Fig. S2 and S3). When we manually corrected all remaining errors, we found that estimates of MPD did not improve (SI Appendix, Fig. S4 and S5). Thus, uncorrected errors do not explain why DADA2 and Deblur estimate MPD poorly.

**FIG. 6:**
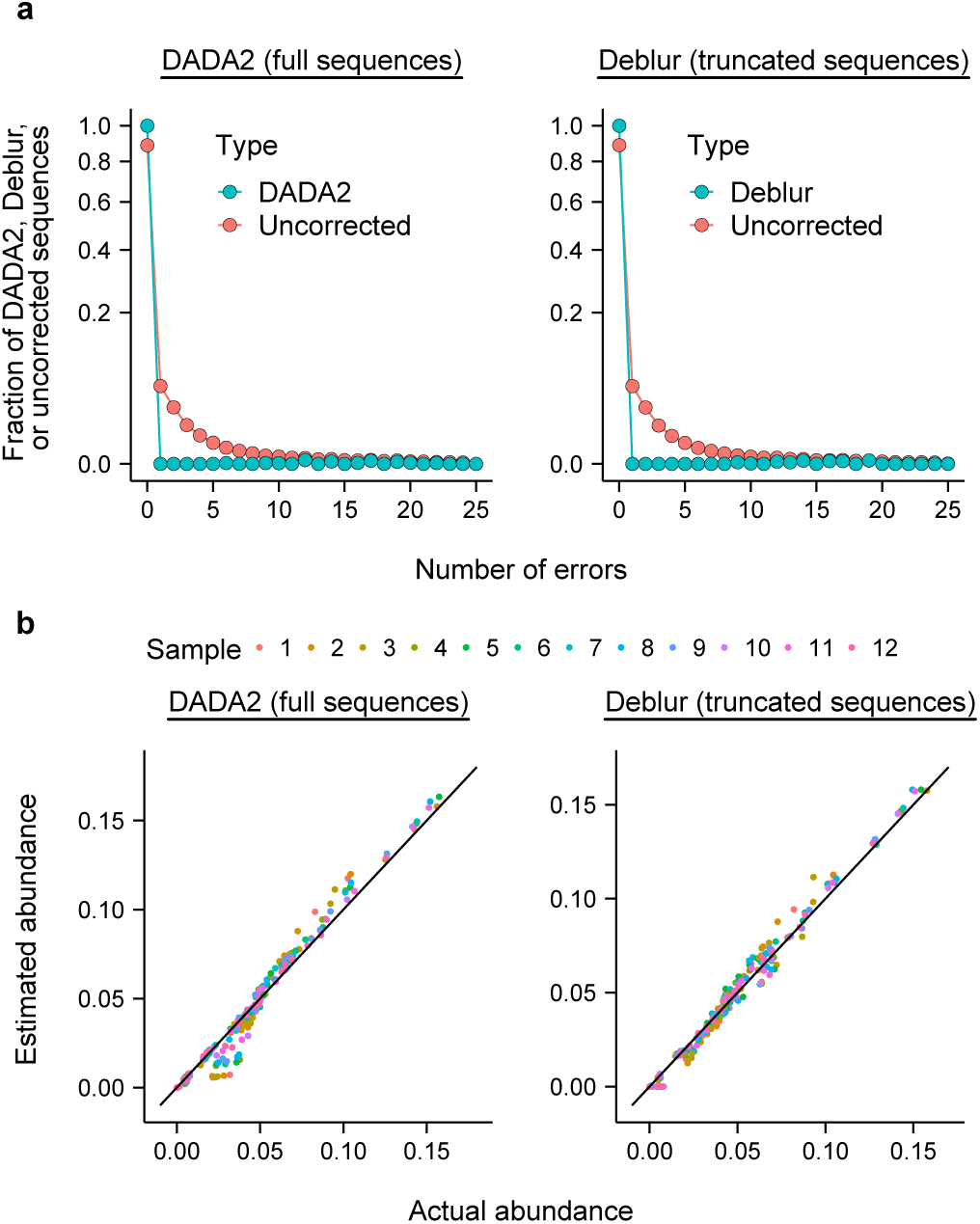
Analysis of sequences outputted by DADA2 and Deblur. (a) Frequency of errors. Values for using no correction for sequencing errors are shown for comparison. (b) Abundance of sequences outputted by Deblur vs. actual abundance. Values shown are for the V4 region of 16S rDNA of an artificial bacterial community. Other regions and sequence types are shown in SI Appendix, Fig. S2, S3, S6, and S7.

Though they had few errors, the sequences outputted by DADA2 and Deblur had abundances different from the sequences in the original sample. Most often, DADA2 and Deblur tools 1) underestimated rare sequences and 2) overestimated sequences at medium or high abundance (Fig. 6, SI Appendix, Fig. S6, and S7). Some rare sequences were missing entirely. Thus, DADA2 and Deblur distorted abundances of sequences. This distortion, rather than uncorrected errors, explains why the tools estimated MPD poorly (see SI Appendix, Fig. S4 and S5).

Past evaluations examined how well DADA2 and Deblur estimated richness [8–10]. In our evaluation, DADA2 and Deblur generally underestimated this measure of alpha diversity (SI Appendix, Fig. S8 and S9). This was expected because some rare sequences were missing from their output. However, the estimates are good compared to leaving sequencing errors uncorrected.

Our evaluation has focused on rDNA sequences from microbial communities. In principle, however, Distanced can be applied to any type of sequence. DADA2 and Deblur use parameters calibrated with data from artificial microbial communities [8, 9]. Their use may be restricted to these data. In contrast, Distanced has no such parameters (see Materials and Methods). We thus evaluated Distanced with antibody sequences, which are highly diverse [16]. Its performance was similar for antibodies as for microbes (Fig. 3 and 5), confirming that it can be applied to a wide range of data. DADA2 could also be evaluated with the antibody sequences, and its performance was poor. Deblur could not be evaluated with this type of sequence (see Materials and Methods).

## Discussion

Though introduced in medicine, the principle of *primum non nocere* (do not harm) should apply to all arenas of science. Towards this end, tools for correcting ribosomal sequences from microbial communities should improve, not worsen, estimates of microbial diversity. We show, unfortunately, that this principle is broken with two popular tools (DADA2 and Deblur). The original sequencing data (with sequencing errors) generally produced better estimates of alpha diversity than did the output of the tools.

The problem has been overlooked by evaluating these tools with a simple measure of diversity (richness) [8–10]. It becomes apparent only when using a more complex measure (MPD) that accounts for sequence abundance and relatedness. Past evaluations had shown that tools distort abundance of sequences [9]. In retrospect, it is unsurprising that the existing tools might estimate MPD poorly.

Our tool (Distanced) does not estimate diversity perfectly, but it does reduce error markedly and consistently. In a display of its flexibility, it estimates diversity for anti-body sequences as accurately as for microbial sequences. No adjustments to the tool were required to accommodate different sequences.

The communities we used to evaluate our tool are artificial, and real microbial communities are more diverse [1]. Our tool has no parameters and was not calibrated using artificial communities. Thus, we expect that it would perform as well on real microbial communities as artificial ones, despite the differences between these communities. Existing tools (DADA2 and Deblur) were calibrated using artificial communities [8, 9]. Their performance with real communities may be even worse than suggested by this evaluation.

DADA2 and Deblur output sequences containing few errors. They are useful tools when the goal of analyzing rDNA sequences is to correct or remove erroneous sequences. However, they estimate alpha diversity poorly when accounting for sequence abundance and relatedness. By using a novel approach, Distanced estimates alpha diversity accurately. With accurate estimates in hand, investigators can answer important questions about microbial diversity. In particular, they can better answer how loss in microbial diversity may affect human health or ecosystem function.

## Materials and Methods

Our method for correcting distances consists of a single equation, and it is derived using Bayess theorem (see SI Appendix, Supporting Information Materials and Methods). It estimates the original distance between two sequences (before introduction of sequencing errors). It requires only 1) the observed distance (after introduction of sequencing errors) and 2) error rates (estimated from quality scores). The equation is

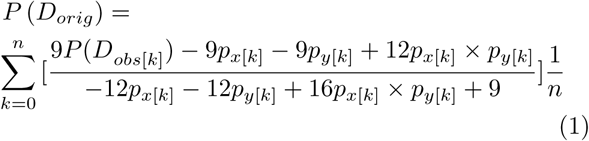

where *P* (*D_orig_*) is the estimated original distance, *P* (*D*_*obs*[*k*]_) is the observed distance at nucleotide position *k*, *p*_*x*[*k*]_ is the sequencing error rate for the first sequence at *k*, and *p*_*y*[*k*]_ is the error rate for the second sequence at *k*, and *n* is the number of nucleotides in the aligned sequences.

The estimate given by eq. [1] is that of the *p* distance, or the fraction of nucleotide letters that differ [17]. This is the type of distance reported in the main text. We also calculated the Jukes-Cantor distance [17], defined as

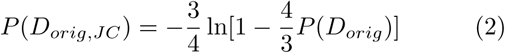

This is a better estimate of the evolutionary distance, which is the number of nucleotide substitutions per site. However, it is undefined when *P* (*D_orig_*) ≥ 0.75, as was the case for some pairs of sequences for some samples. Using either the *p* distance or Jukes-Cantor distance gave similar results when they could be compared (SI Appendix, Fig. S10).

### Simulated reads

We first applied our method to sets of simulated reads. Twenty-five thousand pairs of reads were simulated with *n* = 300 positions. Letters (A, T, C, G) were chosen randomly for one member of the pair. Letters for the other member were chosen to match at a specified distance (e.g., 0.05). Errors were introduced at a rate of 0.0025, which is a typical value for real reads (see SI Appendix, Dataset S9). Errors were introduced under the assumptions they 1) occur independently and 2) are substitutions.

### Real reads

We next applied our method to samples of real (biological) reads. Reads corresponded to three different types of sequences: 16S rDNA of 21 bacterial strains [18], 18S rDNA of 9 fungal strains [19], and synthetic mRNA of 16 different antibodies [20]. The antibody sequences were based on the immunoglobulin G heavy chain of the mouse. Samples are fully described in SI Appendix, Dataset S1. Reference sequences (the actual sequences) were obtained from the publications or, in the case of 16S rDNA, downloaded from https://www.mothur.org/MiSeqDevelopmentData.html.

Reads were minimally processed before analysis with Distanced, DADA2, and Deblur (Fig. 7). Reads retained at each step of processing are reported in SI Appendix, Dataset S2.

**FIG. 7:**
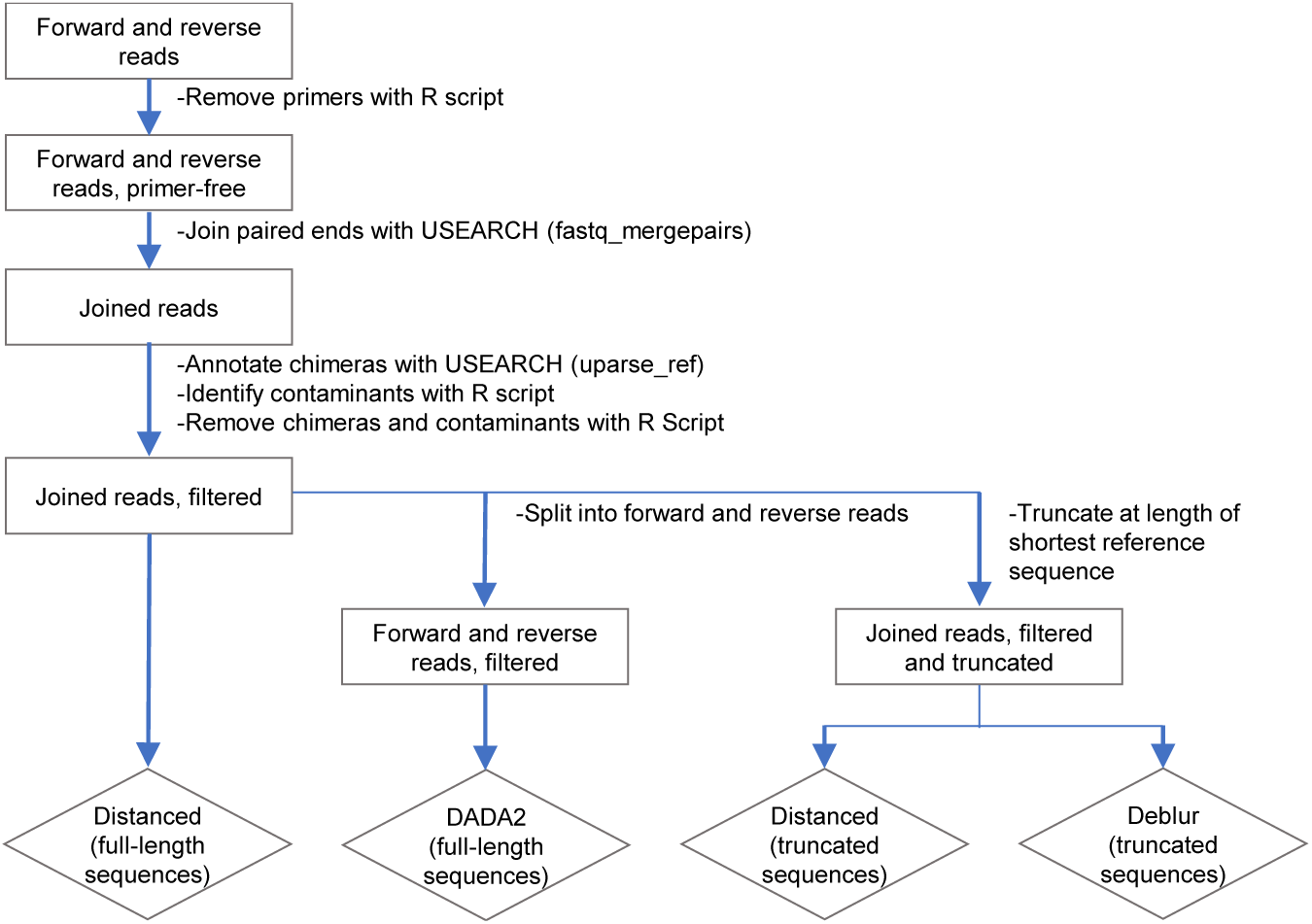
Overview of processing sequence reads for Distanced, DADA2, and Deblur.

Primers were removed using a custom R script. This script enabled removal of primers at both 5 and 3 ends. Primers were present at the 3 end of many ITS1 reads because the original sequences (amplicons) were short, and the read could extend to the very end. The script was not applied to 16S rDNA because primers had already been removed by the authors.

The paired ends of forward and reverse reads were joined with USEARCH (v10.0.240_win32) using parameters in SI Appendix, Dataset S3 and a custom script. For 16S rDNA of bacteria and 18S rDNA, this step served to separate the different regions analyzed.

Reads were annotated as PCR chimeras (concatenations of two parent sequences) using the uparse_ref_command. This command compares reads to both reference sequences and chimera models. The read was annotated as a chimera if it was at least three letters more similar to a chimera model than a reference sequence [21, 22].

Using the output from the uparse_ref command, we annotated contaminants as reads with ≥ 25 differences from reference sequences. Contaminants may originate from the environment, or they may originate from other samples (due to faulty demultiplexing) [23]. The threshold of 25 differences was set to a high value to avoid removing too many good reads (non-contaminants). The high value was needed because the number errors follows a distribution with a long tail (see Fig. 6 and SI Appendix, Fig. S2 and S3). In total, 0.77% of 16S rDNA, 0.005% of 18S rDNA, and 0.03% of antibody mRNA reads were annotated as contaminants. After their annotation, chimeras and contaminants were removed, giving joined and filtered reads.

For DADA2, joined reads were split into individual forward and reverse reads. The splitting was done by matching the ID of the joined and filtered reads with the IDs of the original forward and reverse reads. This step was needed because DADA2 corrects (denoises) sequences prior to joining them. DADA2 (v. 1.8) was subsequently run using these reads, parameters in SI Appendix, Dataset S4, and a custom script. The number of reads inputted, outputted, and remaining at different steps in DADA2 is reported in SI Appendix, Dataset S5.

For Deblur, joined reads were truncated (trimmed) at the 3 end to the length of the shortest reference sequence. This step was needed because Deblur requires reads to be the same length. Deblur was run within QIIME2 (https://qiime2.org) using these reads, parameters in SI Appendix, Dataset S6, and a custom script. The number of reads inputted, outputted, and remaining at different steps in Deblur is reported in SI Appendix, Dataset S7.

Deblur includes a positive filtering step, which removes sequence reads that do not match a reference database. For 16S rDNA, the database was 88% OTUs from GreengeneS13 8 (the default). For 18S rDNA, the database was from UNITE [24]. It was the QIIME release, version 7.2, and with a dynamic threshold value (https://doi.org/10.15156/BIO/587481). For antibody mRNA, we made a reference database containing the original 16 synthetic mRNA sequences [20]. With antibody mRNA, Deblur failed with an error message reported in SI Appendix, Supporting Information Materials and Methods. Thus, Deblur could not be used to analyze antibody sequences.

Distanced was run using a custom R script. For each sample, 1000 reads were randomly subsampled (out of the total number of reads reported in SI Appendix, Dataset S2). If present in a read, ambiguous letters (N) were replaced with an A, T, C, or G (chosen randomly). Reads were aligned against reference sequences with CLUSTAL OMEGA [25, 26]. A matrix of estimated distances was constructed by calculating distances between each pair of sequence reads. Distances were calculated according to eq. [1], observed identities, and instrument-reported error rates. The observed distance was 0 if letters of the nucleotide pair matched and 1 if they did not. Gaps were considered mismatches (observed distance = 1) if they appeared in only one sequence of the pair; otherwise, they were ignored.

Mean pairwise distance (MPD) was the mean of distances in the matrix. The diagonal elements in the matrix were not included. Values of MPD were also calculated for sequences outputted by DADA2, sequences outputted by Deblur, and for sequences with errors left uncorrected (joined and filtered reads).

This subsampling of 1000 reads and calculation of MPD was iterated 100 times per sample. Reported values of MPD and other variables are means of these 100 iterations.

Distanced was run with both truncated and nontruncated (full-length) sequences. This enabled separate comparison to Deblur and DADA2. For the V34 region of 16S rDNA, DADA2 outputted fewer than 1000 reads for some samples (see SI Appendix, Dataset S5). For the full-length sequences for this region, 900 reads were thus subsampled.

A matrix of actual distances was determined by 1) finding a matching reference sequence for each read and 2) calculating the distance between these matches. The matching reference sequence was that with highest identity with the read. Actual MPD was calculated from this matrix. Numerical values for MPD (as well as richness) are in SI Appendix, Dataset S8.

Instrument-reported error rates were calculated from quality scores (*Q*) as 10^−*Q/*10^. Gaps had no quality scores and were assigned an error rate of 0. Actual error rates were calculated by comparing uncorrected and actual reads. Numerical values for error rates are in SI Appendix, Dataset S9.

### Data sharing

All scripts are available at https://github.com/thackmann/Distanced. All sequence data is available at sources indicated by SI Appendix, Dataset S1.

## SI Appendix

### Derivation of eq. [1] in main text

We will derive an equation that estimates the distance between two nucleic acid sequences before introduction of sequencing errors. The derivation will use Bayes’ theorem.

Let *X* and *Y* be the letters of two sequences (Fig. S11). *X*_[*k*]_ and *Y*_[*k*]_ refer to a letter at a given nucleotide position *k* within the sequences, and sequences have a total of *n* positions. Before sequencing (introduction of errors), the letters were originally *X_orig_* and *Y_orig_*. After sequencing (introduction of errors), some letters change, and the measured sets of letters become *X_obs_* and *Y_obs_*. Let *D_orig_* be letters in *X_orig_* and *Y_orig_* that are different (when compared at a given position *k*), and *D_obs_* be letters different between *X_obs_* and *Y_obs_*. The letters that are identical are *I_orig_* and *I_obs_*. We partition *D_obs_* as 1) *D_obs_*1, which originate from *D_orig_*, and 2) *D*_*obs*2_, which originate from *I_orig_*.

Our goal is to calculate the original distance, *P* (*D_orig_*), or distance before introduction of errors. It is defined as *P* (*D_orig_*) = *n_Dorig_/n*, where *n_Dorig_* is the number of positions in *D_orig_*. We will calculate it from 1) the observed distance, *P* (*D_obs_*), or distance after introduction of errors, and 2) the error rates *p_x_* and *p_y_* (defined below).

#### Distance at a given position

To estimate the distance between *X* and *Y* in total, we will first estimate the distance at a given position in *X* and *Y*. *P* (*D*_*obs*[*k*]_) is the observed distance at position *k* and is either 1 (letters different) or 0 (letters identical). Following Fig. S12, we can partition it as

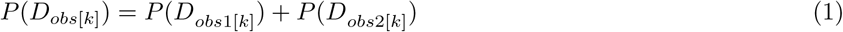

By Bayes’ theorem

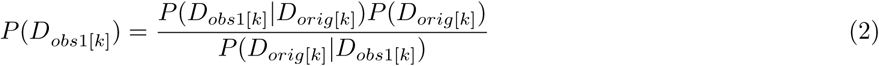

and

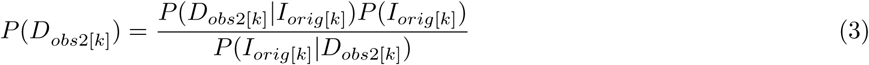

We note *P* (*D*_*orig*[*k*]_) = 1 − *P* (*I*_*orig*[*k*]_), substitute eq. [2] and [3] into [1], and solve for *P* (*D*_*orig*[*k*]_) to give

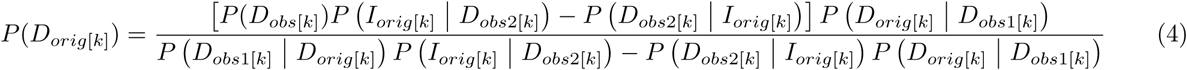

Next, we find expressions for *P* (*D*_*orig*[*k*]_ | *D*_*obs*1[*k*]_), *P* (*I*_*orig*[*k*]_ | *D*_*obs*2[*k*]_), *P* (*D*_*obs*1[*k*]_ | *D*_*orig*[*k*]_), and *P* (*D*_*obs*2[*k*]_ | *I*_*orig*[*k*]_. Because all *D*_*obs*1[*k*]_ originate from *D*_*orig*[*k*]_ and all *D*_*obs*2[*k*]_ originate from *I*_*orig*[*k*]_ (Fig. S11),

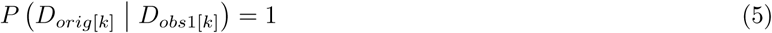

and

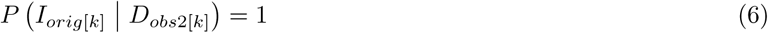

We partition *P* (*D*_*obs*1[*k*]_ | *D*_*orig*[*k*]_) and *P* (*D*_*obs*2[*k*]_ | *I*_*orig*[*k*]_) as

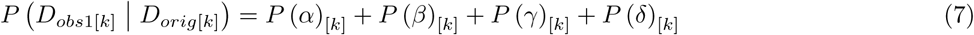

and

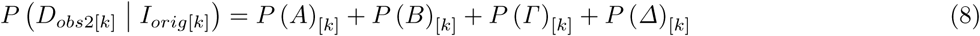

with terms defined in Table S1. For example, *P* (*α*)_[*k*]_ = (1 − *p*_*x*[*k*]_)(1 − *p*_*y*[*k*]_) is the probability that neither *X*_[*k*]_ nor *Y*_[*k*]_ change (no errors were introduced) after sequencing, given the letters were different before sequencing (i.e., *X*_[*k*]_ and *Y*_[*k*]_ belong to *D_orig_*). The terms *p*_*x*[*k*]_ and *p*_*y*[*k*]_ are probabilities for change (error rates) for *X*_[*k*]_ and *Y*_[*k*]_, respectively. We assume all errors are substitutions (not insertions or deletions), giving one of three equally probable outcomes per position per sequence. Quality scores (*Q*) can be used to calculate the error rates [e.g., *p*_*x*[*k*]_ = 10^−(*Q*_*X*[*k*]_/10)^].

Substituting expressions for Table 3 into eq. [7] and [8] gives

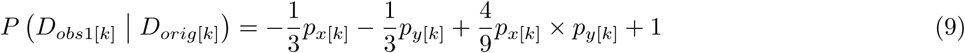

and

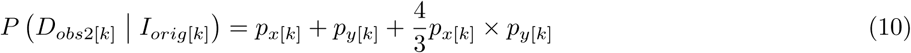

By substituting eq. [5], [6], [9], and [10] into eq. [4], we yield

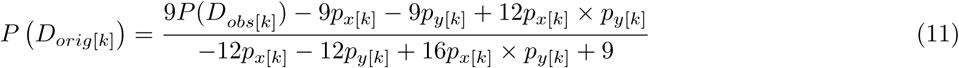

We can derive eq. [11], with the same result, if we follow Fig. S12b and partition as *P* (*I*_*obs*[*k*]_) as *P* (*I*_*obs*[*k*]_) = *P* (*I*_*obs*1[*k*]_) + *P* (*I*_*obs*2[*k*]_) (not shown).

To estimate distance of *X* and *Y* in total, we average *P* (*D*_*orig*[*k*]_) across all *n* positions

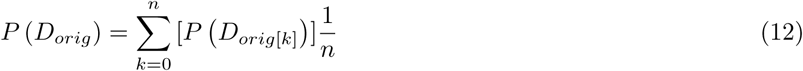

This approach assumes all changes (errors) occur independently (i.e., an error occurring at *k* = 0 does not change the probability of an error at *k* = 1). Eq. [12] can be expanding by substituting in eq. [11], giving

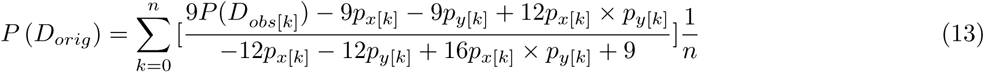

Eq. [13] is eq. [1] in the main text.

#### Failure of Deblur on antibody sequence reads

To analyze antibody sequence reads with Deblur, we made a reference database containing the original 16 synthetic mRNA sequences. However, Deblur failed with the error message:

~~~
 File “/home/qiime2/miniconda/envs/qiime2-2018.6/lib/python3.5/site-packages/deblur/workflow.py”,
line 467, in remove artifacts seqs
 if (float(line[2]) >= sim_thresh) and \
IndexError: list index out of range
~~~

The reason for the failure could not be determined. To troubleshoot, we performed the analysis with other reference databases or sequence reads. For example, we analyzed 1) the antibody sequence reads with a non-antibody reference database (e.g., 88% OTUs from Greengenes 13_8) and 2) non-antibody sequence reads (e.g., 18S rDNA from fungi) with the antibody reference database. Deblur did not fail in these cases, though all reads were removed by the positive filter. This suggests that the sequence reads and reference database for antibodies were formatted correctly, and the cause of Deblur’s failure was more deeply seated.

#### Dataset captions

DatasetS1.xlsx. Description of samples analyzed in this study.

DatasetS2.xlsx. Retention of reads during initial processing.

DatasetS3.xlsx. Parameters for merging reads with USEARCH.

DatasetS4.xlsx. Parameters for correcting sequences with DADA2.

DatasetS5.xlsx. Retention of reads during steps of DADA2.

DatasetS6.xlsx. Parameters for correcting sequences with Deblur.

DatasetS7.xlsx. Retention of reads during steps of Deblur.

DatasetS8.xlsx. Measures of diversity for samples analyzed in this study.

DatasetS9.xlsx. Error rates for samples analyzed in this study.

*Note: Datasets are not included with pre*-*print*.

**FIG. S1:**
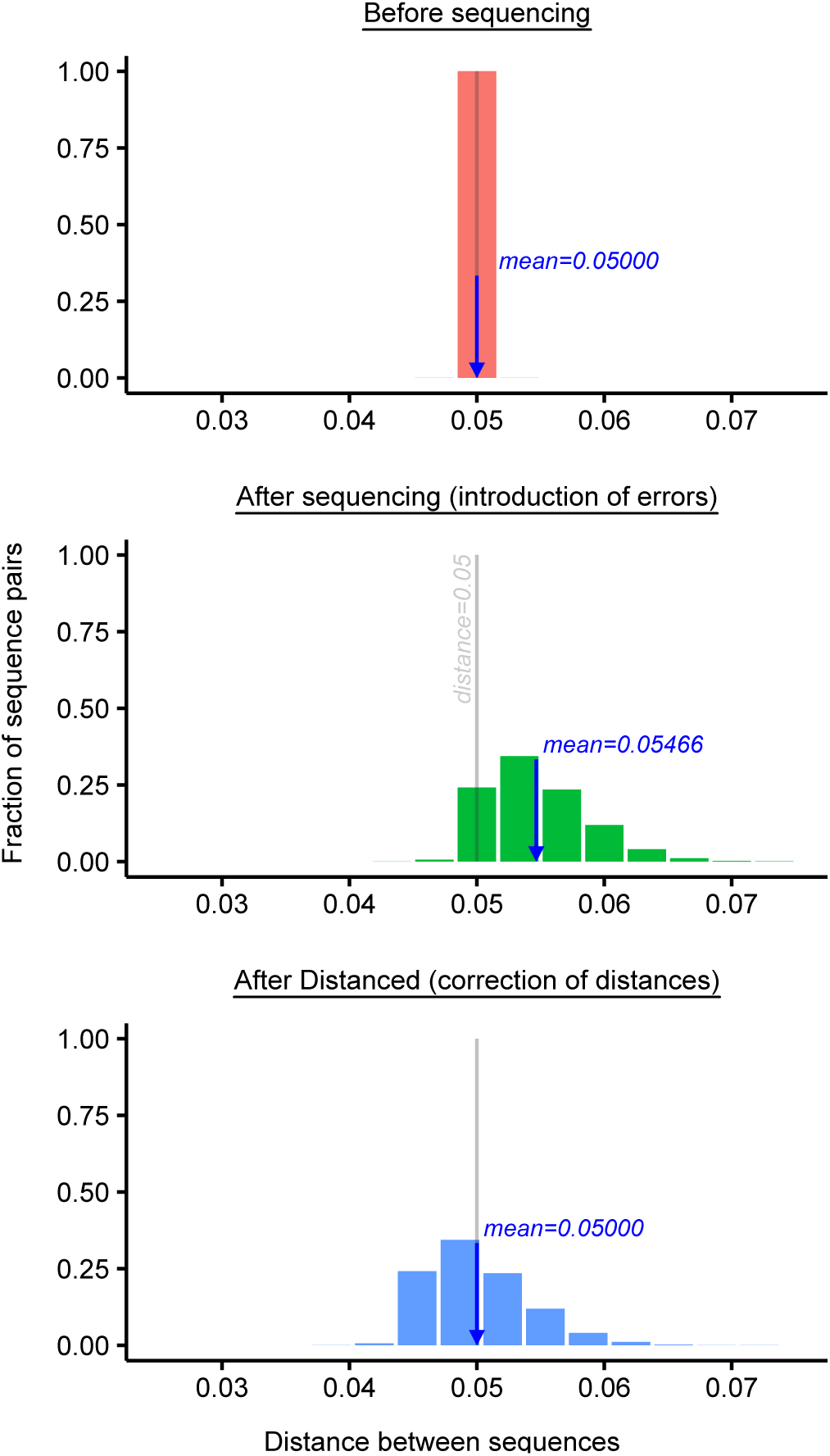
Performance of Distanced in correcting distances when applied to simulated sequence reads. The mean distance after applying Distanced equaled that before sequencing, indicating the correction was accurate. The original distance (0.05) was arbitrary, and the correction was accurate for other values (not shown).

**FIG. S2:**
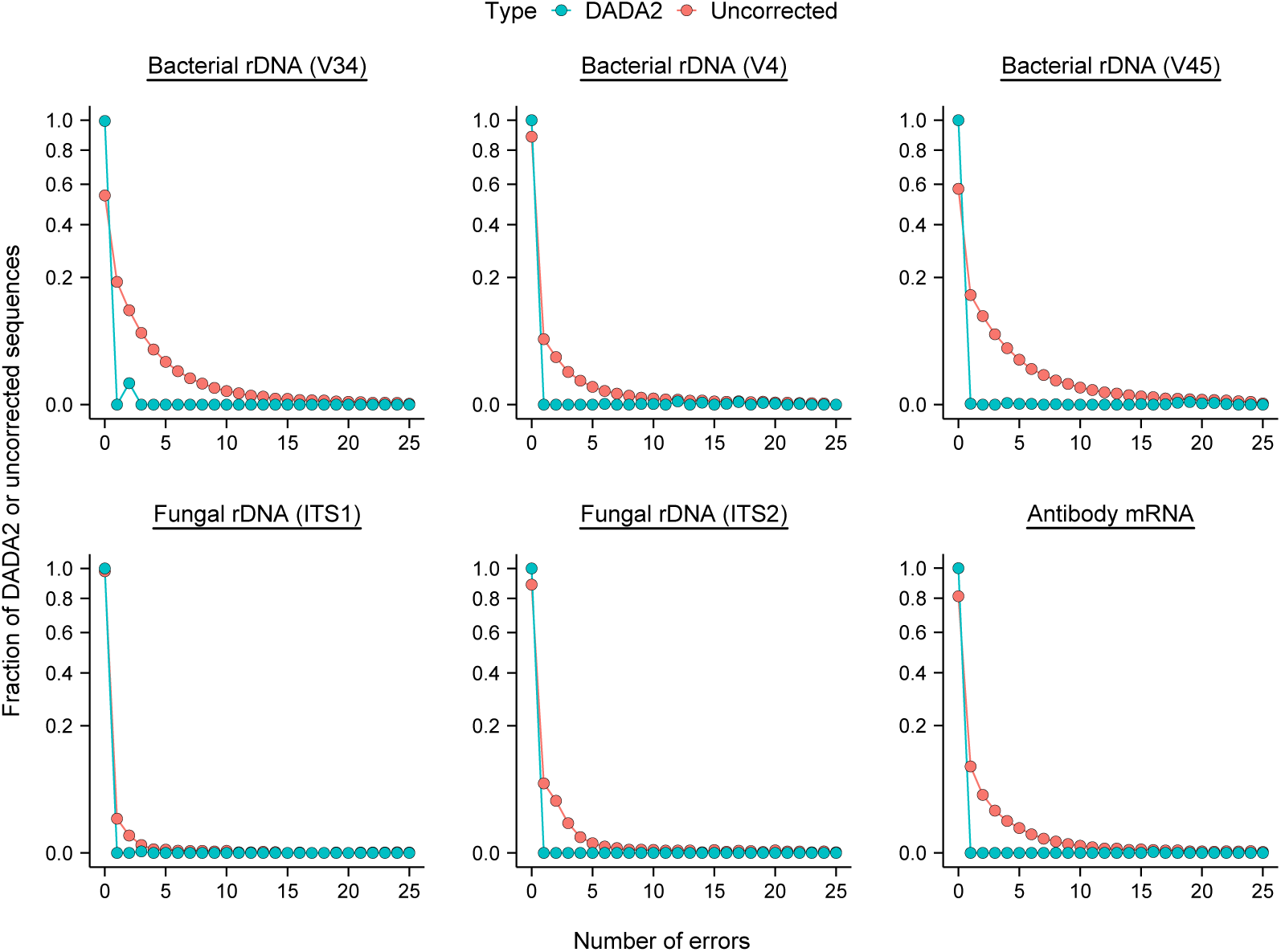
Frequency of errors in sequences outputted by DADA2. Values for when using no correction for sequencing errors are shown for comparison.

**FIG. S3:**
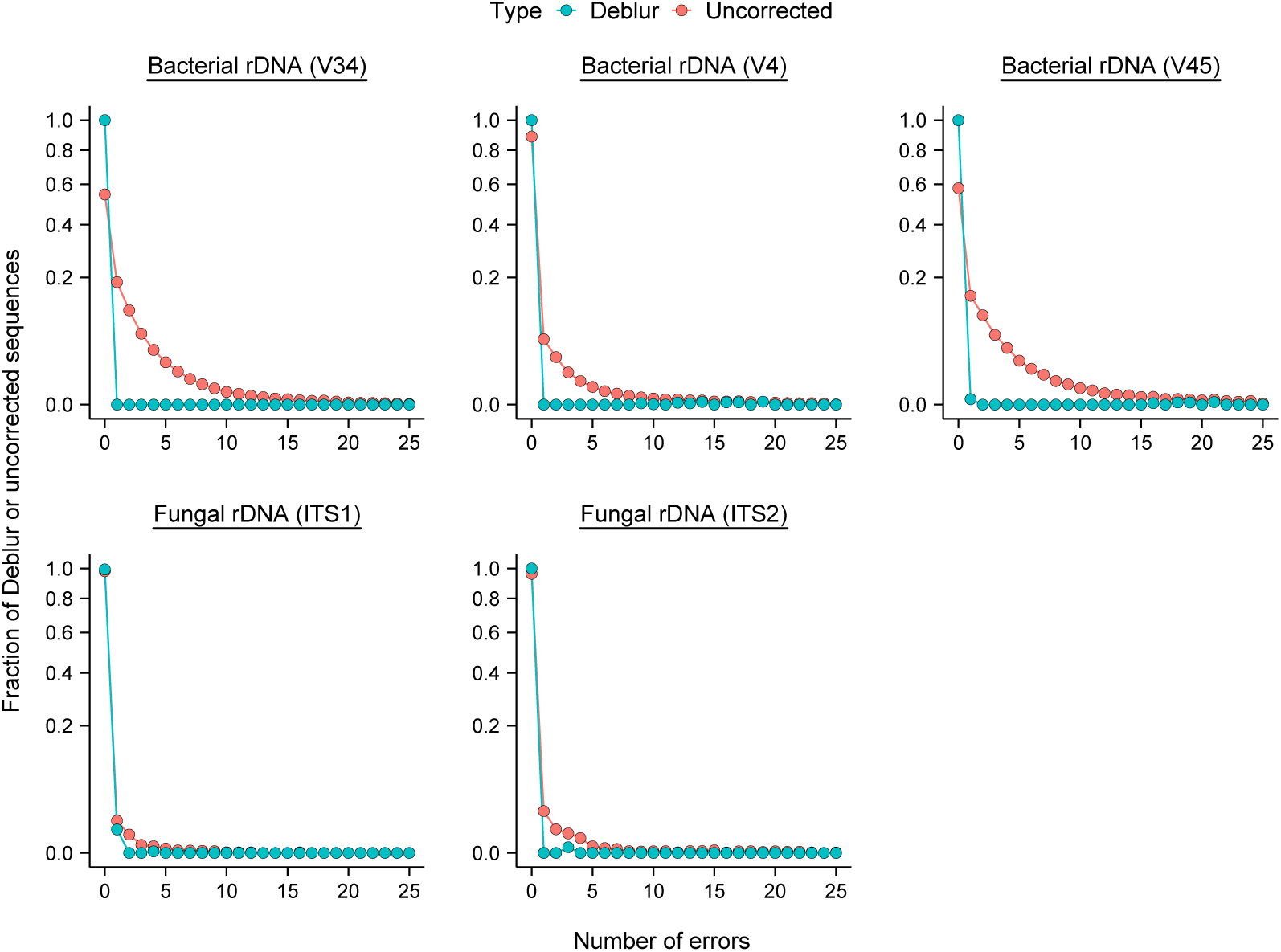
Frequency of errors in sequences outputted by Deblur. Values for when using no correction for sequencing errors are shown for comparison.

**FIG. S4:**
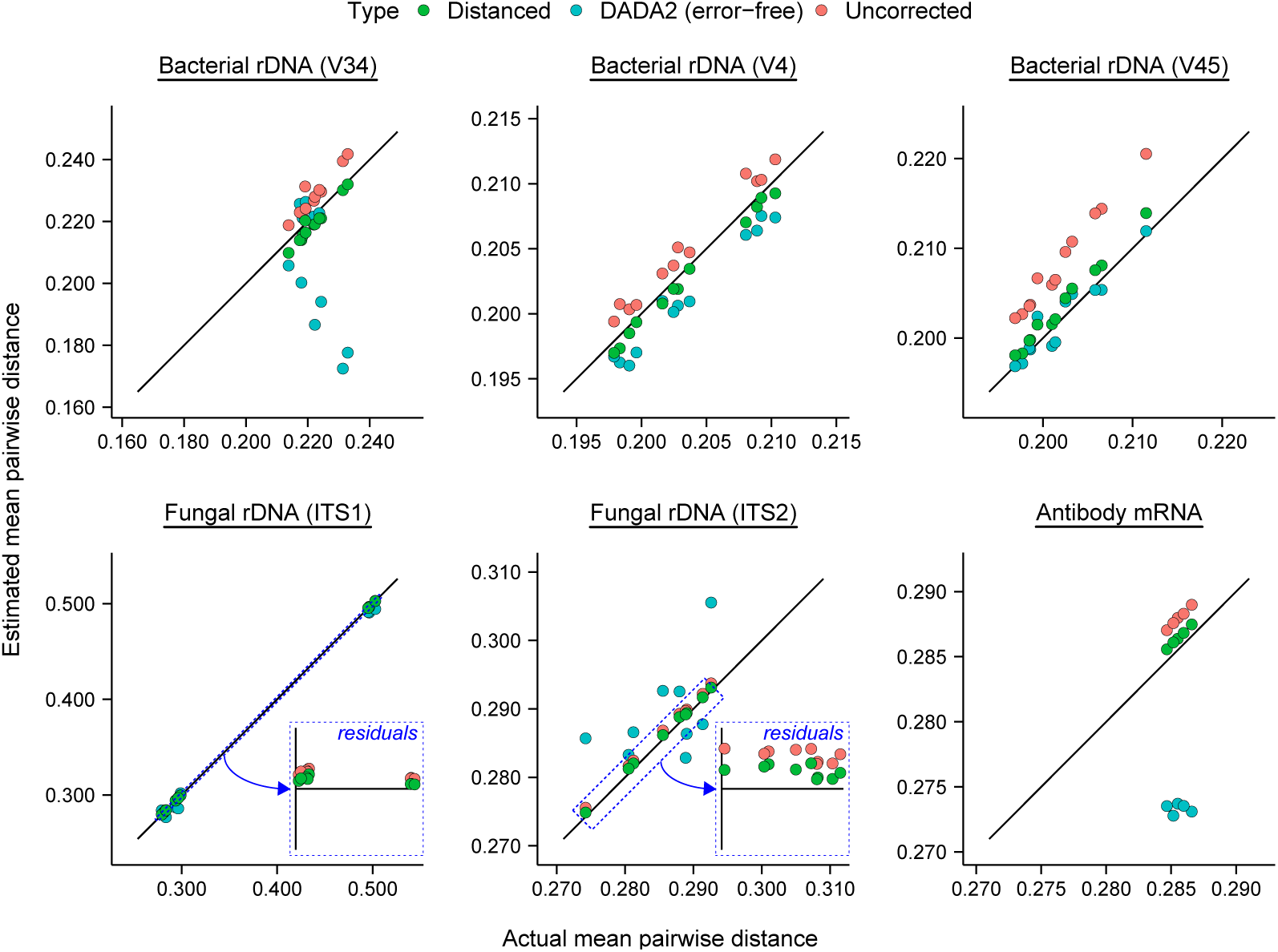
Performance of Distanced vs. DADA2 in estimating alpha diversity (mean pairwise distance) of ribosomal DNA from artificial microbial communities or mRNA of antibodies. Values are as Fig. 3 in the main text, except DADA2 sequences have been manually corrected to remove all remaining errors. Errors were corrected by 1) finding a matching reference sequence for each DADA2 sequence and 2) calculating distance between these matches. After correcting errors, DADA2 sequences differed from the actual sequences only in their abundance. Thus, poor estimates of mean pairwise distance are due to poor estimates of abundance.

**FIG. S5:**
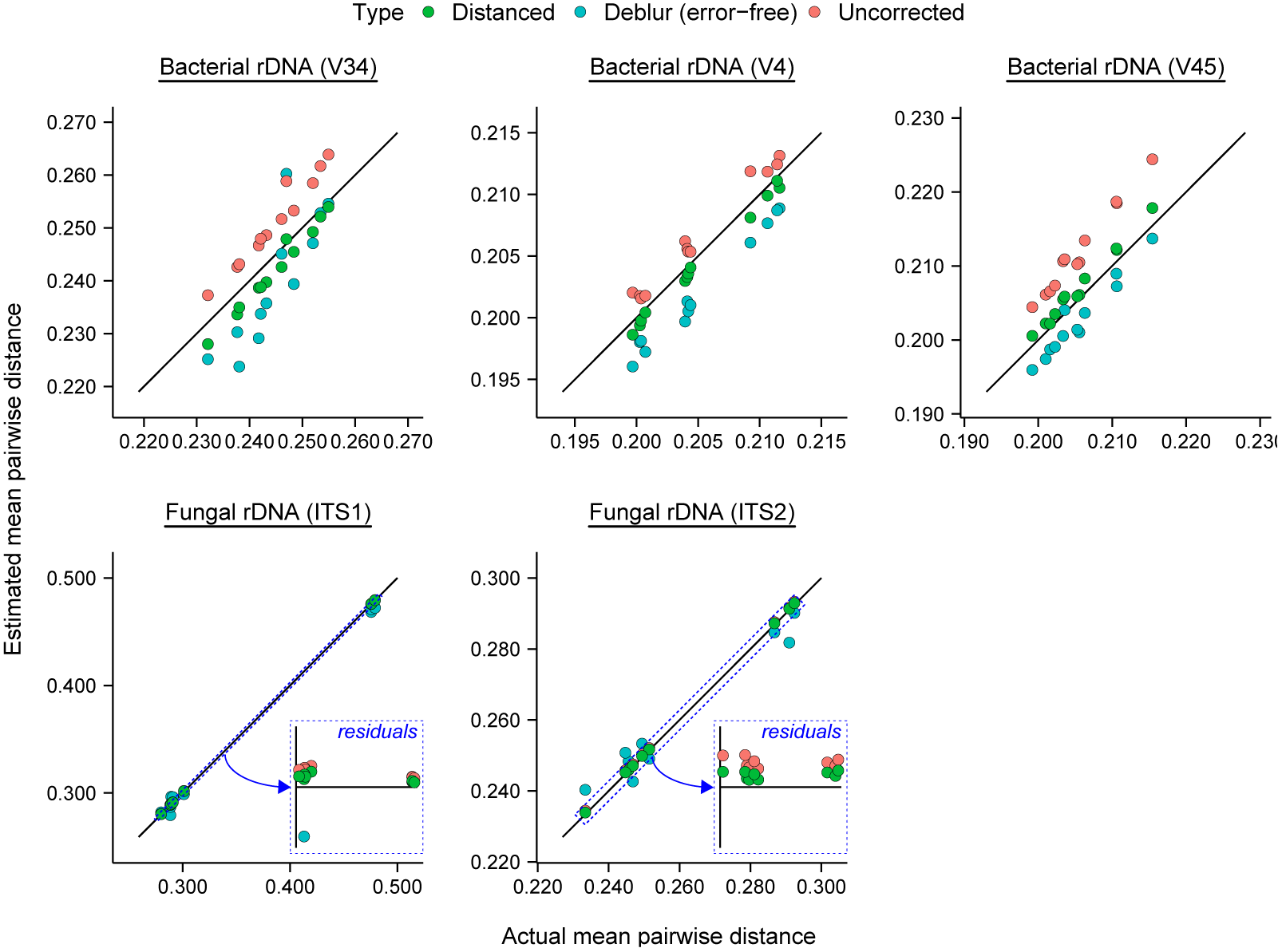
Performance of Distanced vs. Deblur in estimating alpha diversity (mean pairwise distance) of ribosomal DNA from artificial microbial communities. Values are as Fig. 4 in the main text, except Deblur sequences have been manually corrected to remove all remaining errors. Error correction is explained in Fig. S4.

**FIG. S6:**
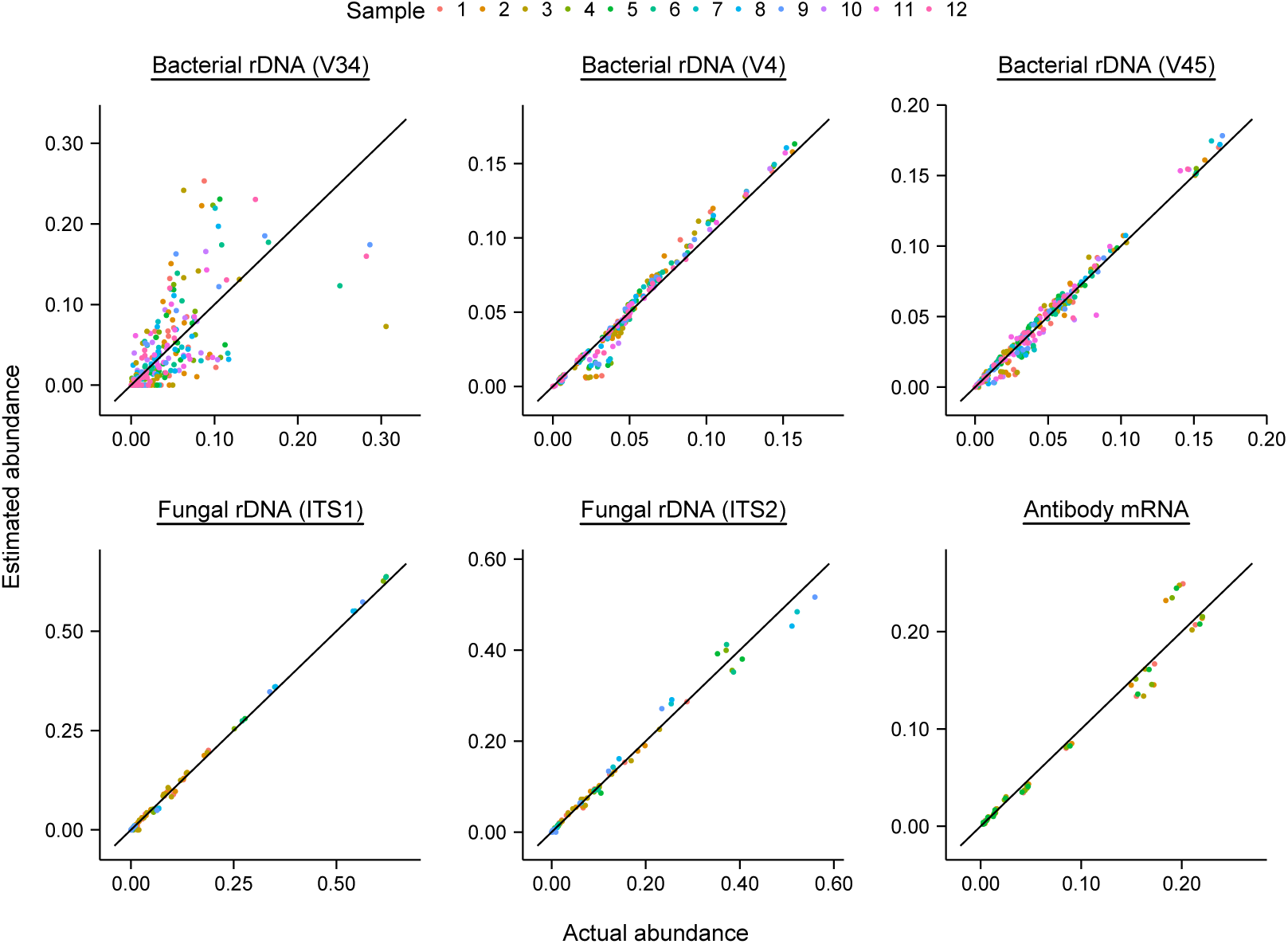
Abundance of sequences outputted by DADA2 vs. actual abundance.

**FIG. S7:**
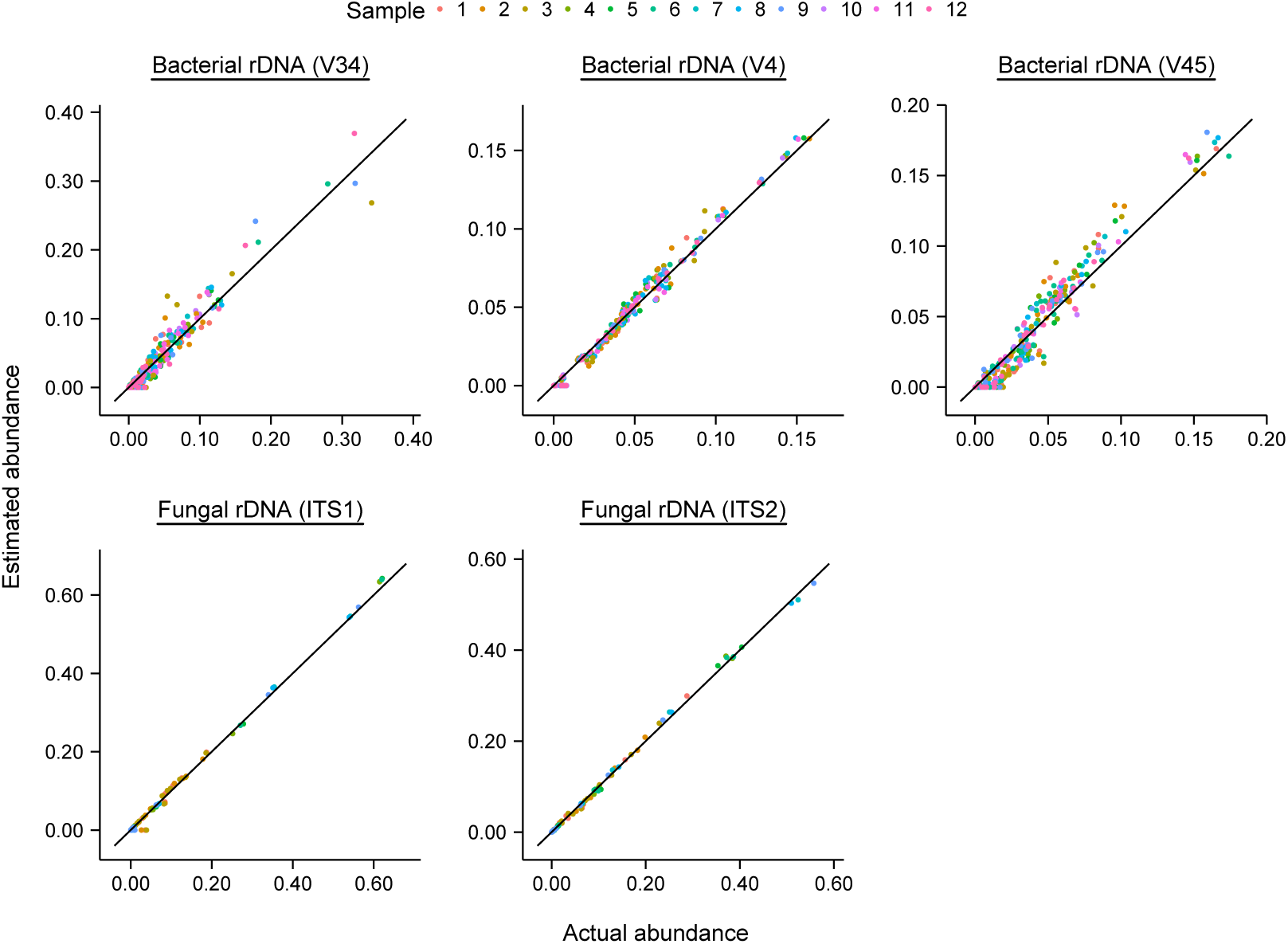
Abundance of sequences outputted by Deblur vs. actual abundance.

**FIG. S8:**
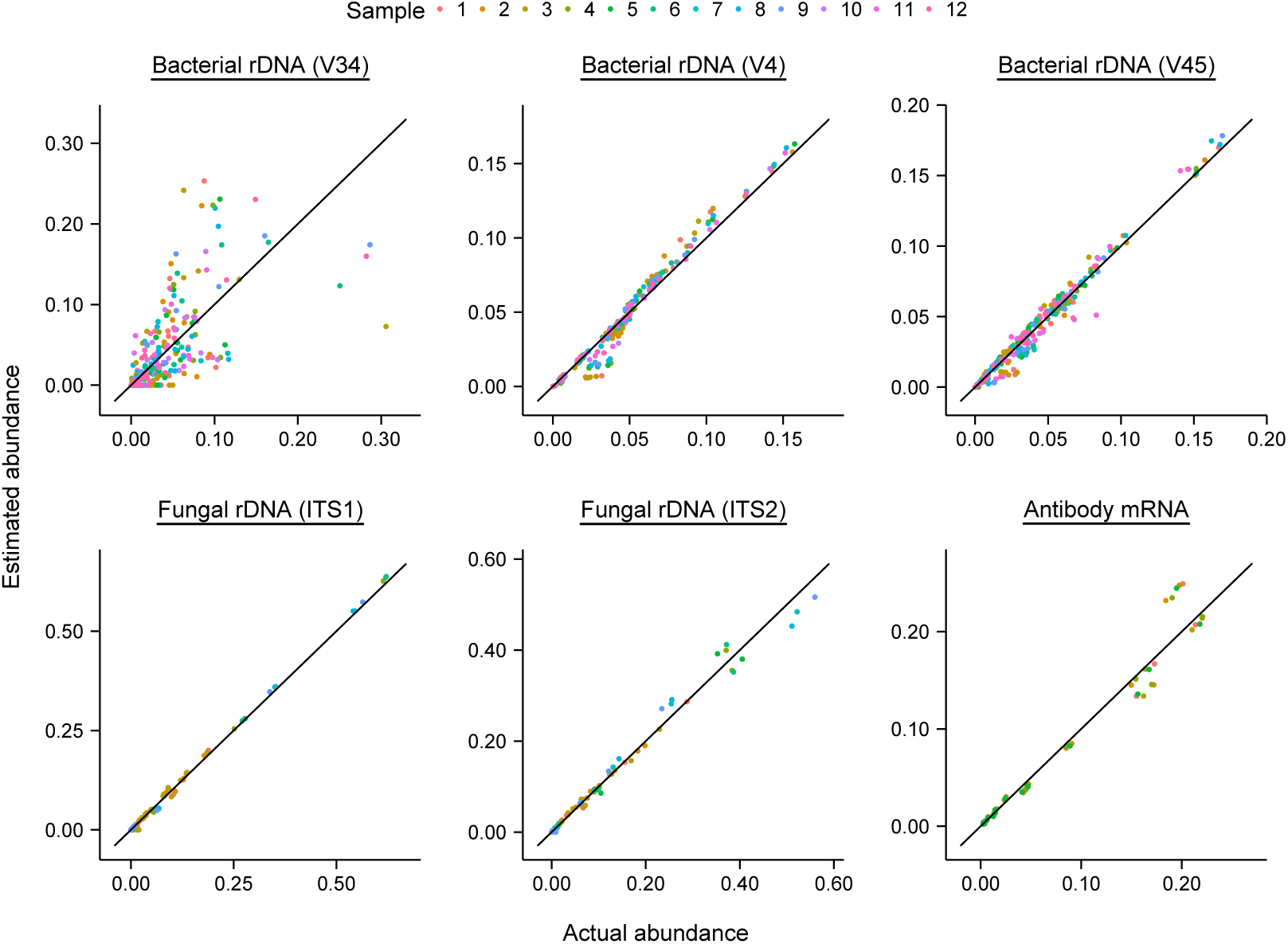
Performance of DADA2 in estimating richness of ribosomal DNA from artificial microbial communities. mRNA from antibody mixtures is included for comparison. Errors in sequence letters were corrected by DADA2. For comparison, we show estimates of richness obtained when either 1) using no correction or 2) manually correcting all remaining errors in DADA2 sequences [DADA2 (error-free)]. Full-length sequences were those analyzed.

**FIG. S9:**
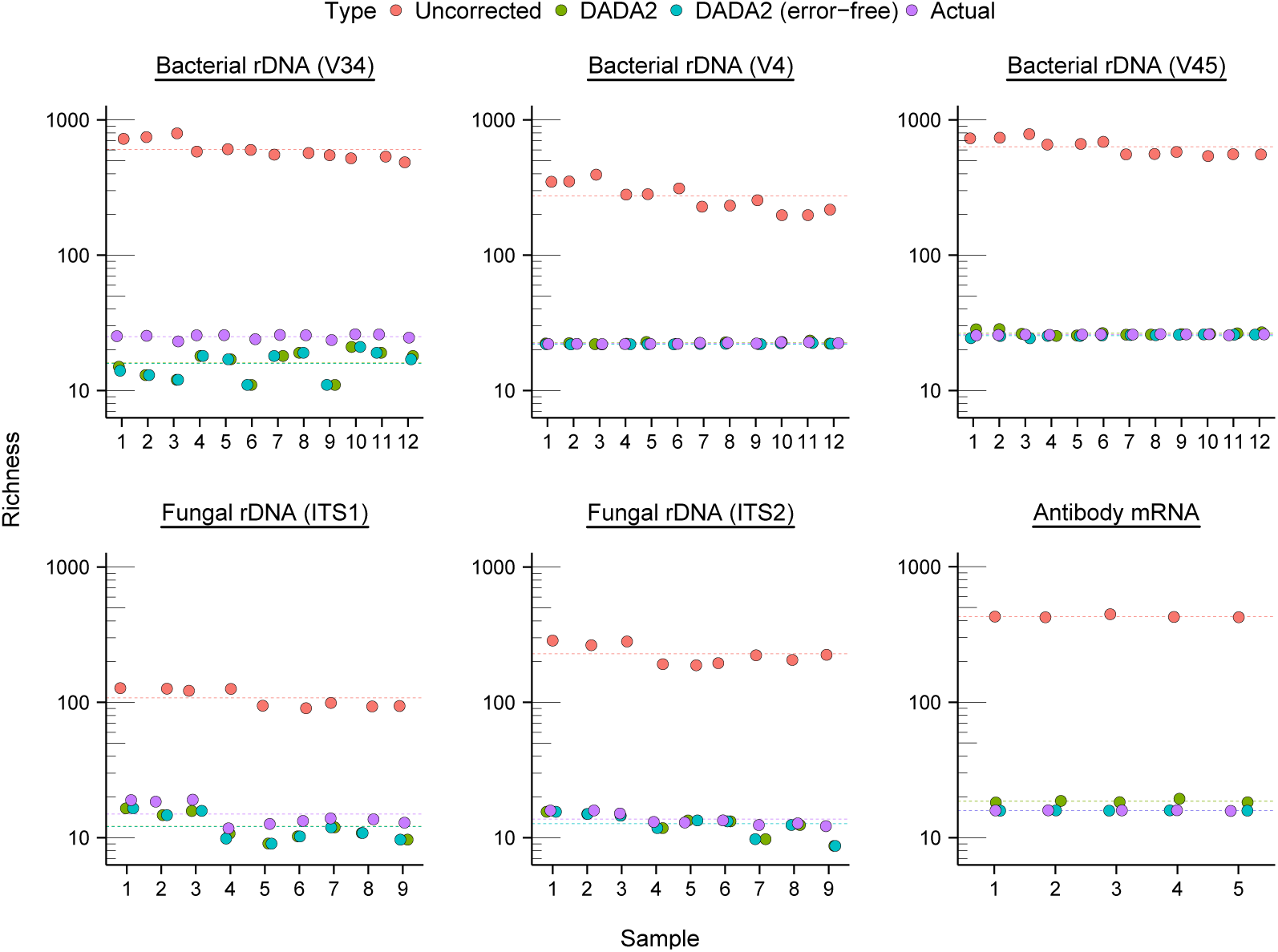
Performance of Deblur in estimating richness of ribosomal DNA from artificial microbial communities. Errors in sequence letters were corrected by Deblur. For comparison, we show estimates of richness obtained when either 1) using no correction or 2) manually correcting all remaining errors in Deblur sequences [Deblur (error-free)]. Full-length sequences were those analyzed.

**FIG. S10:**
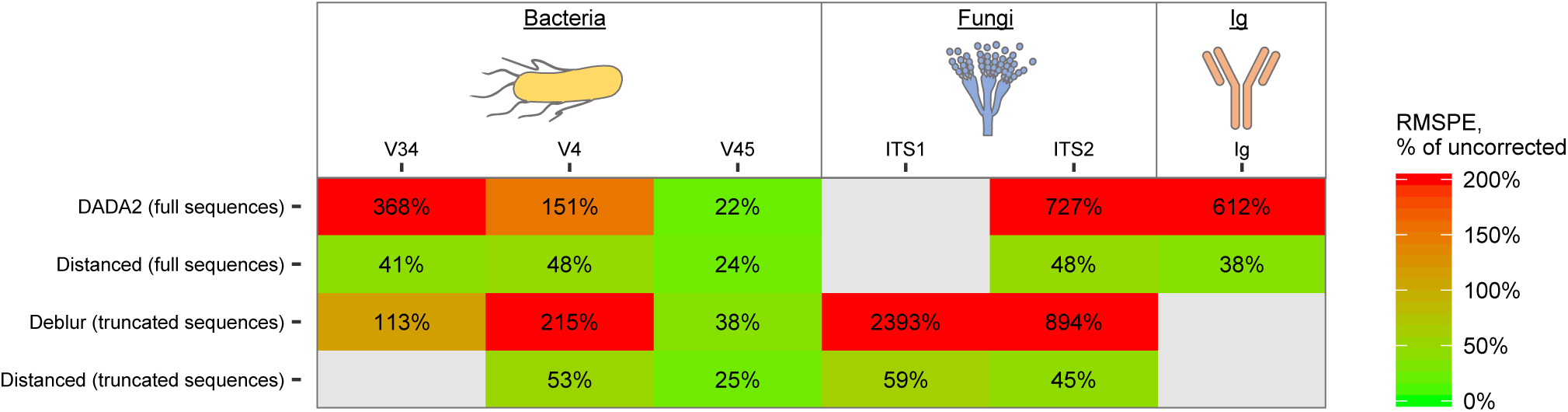
Error in estimating alpha diversity (mean pairwise distance) of artificial microbial communities and antibody mixtures by Distanced, DADA2, and Deblur. The current figure shows errors obtained using Jukes-Cantor distances, whereas Fig. 5 shows those obtained using *p* distances. For certain samples of the V34 and ITS1 regions, the *p* distance ≥ 0.75 for one or more pairs of sequences, and no estimate of the Jukes-Cantor distance could be made.

**FIG. S11:**
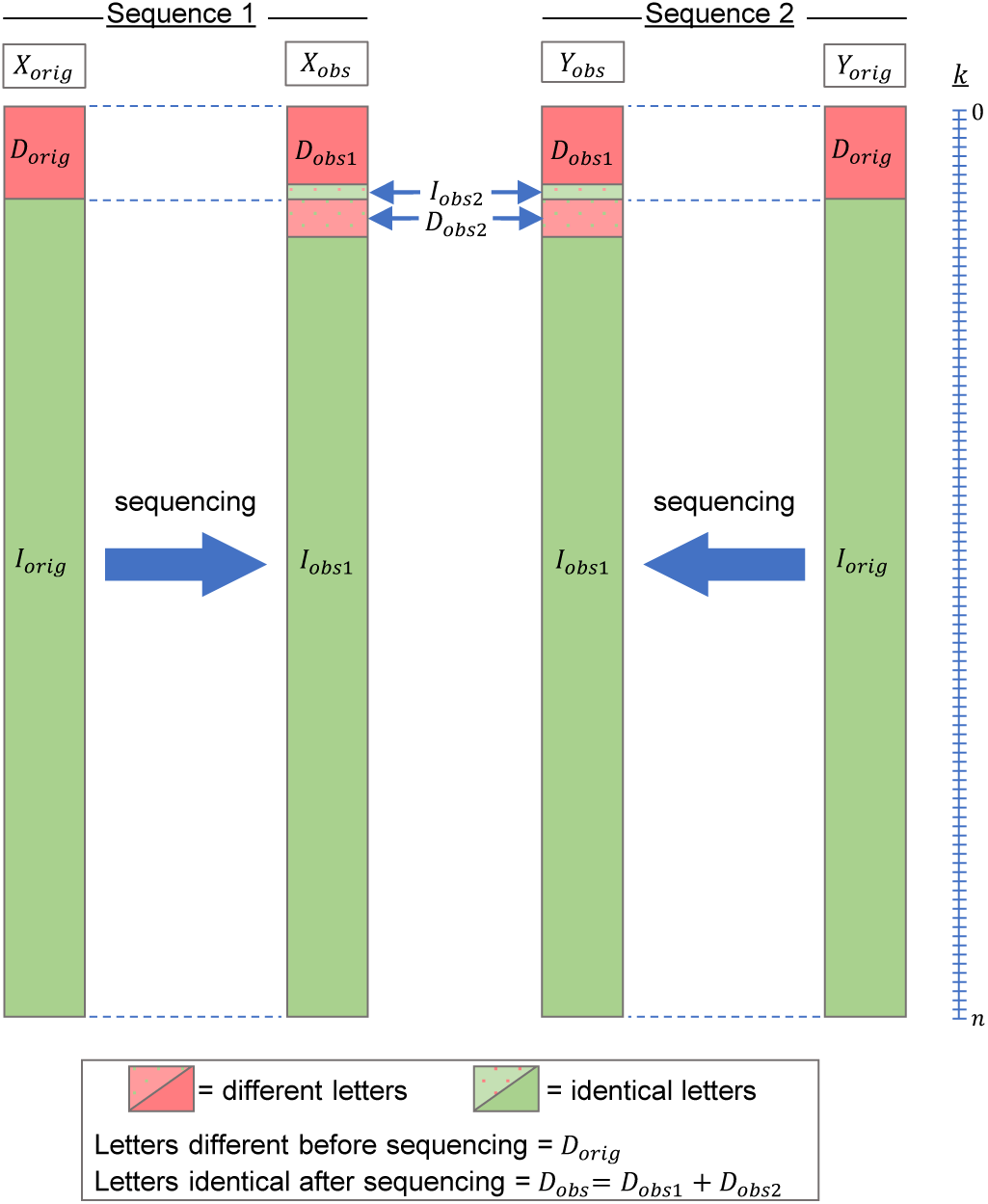
Two DNA sequences before and after sequencing, illustrating of terms in our equations for estimating distances between sequences. Letters in the two sequences are *X* and *Y*. Letters between *X* and *Y* that are different are *D*, and identical letters are *I*. The subscripts *orig* and *obs* (e.g., in *X_orig_* and *X_obs_*) refer to conditions before and after sequencing. Each letter has a position *k*, and there are a total of *n* positions. For illustration, letters in *I_orig_* are grouped separately from letters in *D_orig_*, though they would be interspersed in a real sequence. Letters in *D*_*obs*1_, *D*_*obs*2_, *I*_*obs*1_, and *I*_*obs*2_, are grouped in the same way.

**FIG. S12:**
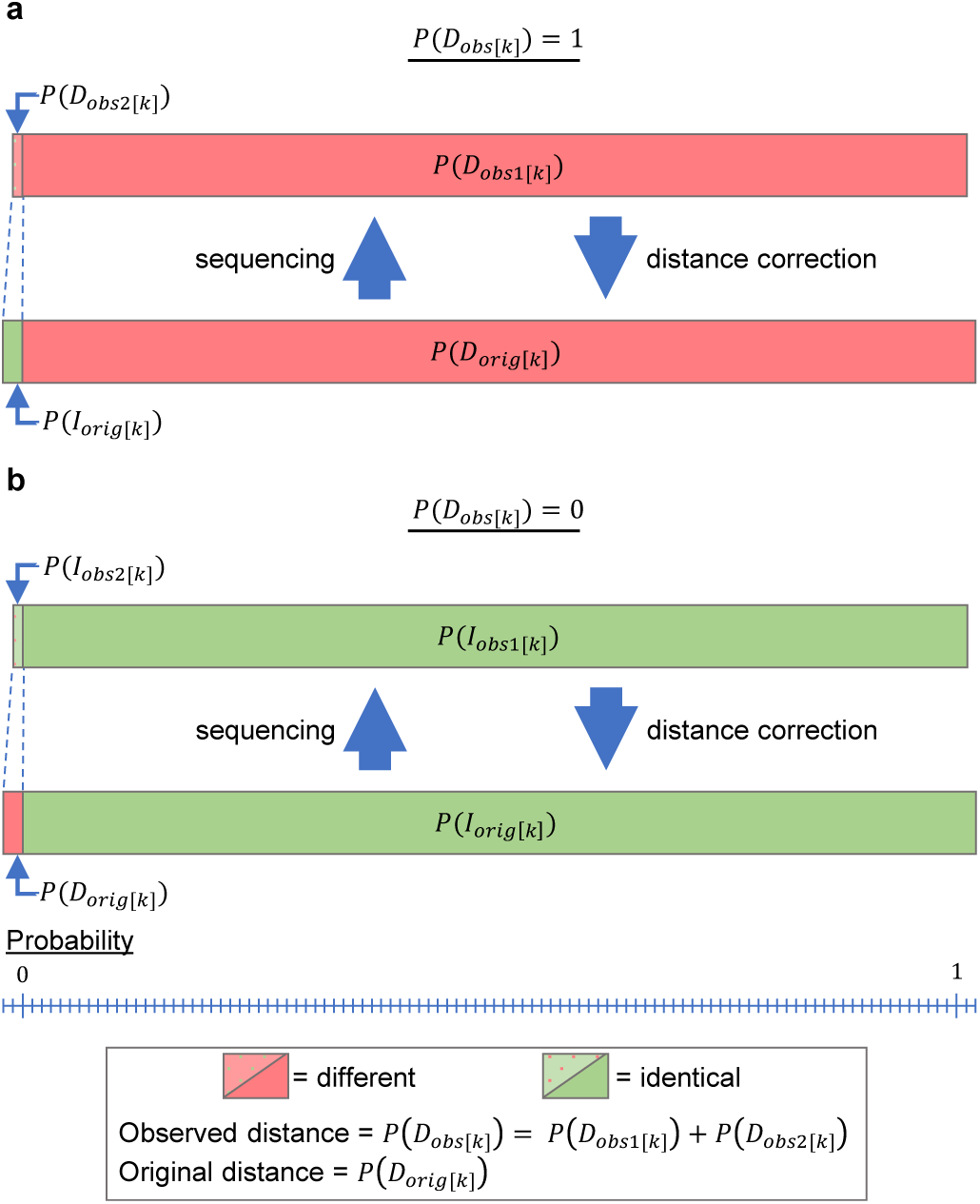
A given sequence position (*k*) before and after sequencing, illustrating of terms in our equations for correcting sequence identity. (a) Condition where letters *X*_*obs*[*k*]_ and *Y*_*obs*[*k*]_ are different [*P* (*D*_*obs*[*k*]_) = 1]. (b) Condition where letters *X*_*obs*[*k*]_ and *Y*_*obs*[*k*]_ are identical [*P* (*D*_*obs*[*k*]_) = 0].

**TABLE I:** Definition of terms in eq. [7] and [8] (and related terms)

| Term<sup>1,2</sup> | Expression | Original (prior) condition | Observed (posterior) condition |  |  |
| --- | --- | --- | --- | --- | --- |
| | | Identical or different? | Identical or different? | Error (change) in $X_{[k]}$ | Error (change) in $Y_{[k]}$ |
| $P(\alpha)_{[k]}$ | $(1 - p_{x[k]}) (1 - p_{y[k]})$ | $X_{[k]}$ and $Y_{[k]}$ different | $X_{[k]}$ and $Y_{[k]}$ different | - | - |
| $P(\beta)_{[k]}$ | $\frac{2}{3} p_{x[k]} (1 - p_{y[k]})$ | | | + | - |
| $P(\gamma)_{[k]}$ | $\frac{2}{3} (1 - p_{x[k]}) p_{y[k]}$ | | | - | + |
| $P(\delta)_{[k]}$ | $\frac{7}{9} p_{x[k]} \times p_{y[k]}$ | | | + | + |
| $P(\varepsilon)_{[k]}$ | 0 | | $X_{[k]}$ and $Y_{[k]}$ identical | - | - |
| $P(\zeta)_{[k]}$ | $\frac{1}{3} p_{x[k]} (1 - p_{y[k]})$ | | | + | - |
| $P(\eta)_{[k]}$ | $\frac{1}{3} (1 - p_{x[k]}) p_{y[k]}$ | | | - | + |
| $P(\theta)_{[k]}$ | $\frac{2}{9} p_{x[k]} \times p_{y[k]}$ | | | + | + |
| $P(A)_{[k]}$ | 0 | $X_{[k]}$ and $Y_{[k]}$ identical | $X_{[k]}$ and $Y_{[k]}$ different | - | - |
| $P(\beta)_{[k]}$ | $p_{x[k]} (1 - p_{y[k]})$ | | | + | - |
| $P(\Gamma)_{[k]}$ | $(1 - p_{x[k]}) p_{y[k]}$ | | | - | + |
| $P(\Delta)_{[k]}$ | $\frac{2}{3} p_{x[k]} \times p_{y[k]}$ | | | + | + |
| $P(E)_{[k]}$ | $(1 - p_{x[k]}) (1 - p_{y[k]})$ | | $X_{[k]}$ and $Y_{[k]}$ identical | - | - |
| $P(Z)_{[k]}$ | 0 | | | + | - |
| $P(H)_{[k]}$ | 0 | | | - | + |
| $P(\Theta)_{[k]}$ | $\frac{1}{3} p_{x[k]} \times p_{y[k]}$ | | | + | + |

<sup>1</sup>As expected, $P(\alpha)_{[k]} + P(\beta)_{[k]} + P(\gamma)_{[k]} + P(\delta)_{[k]} + P(\varepsilon)_{[k]} + P(\zeta)_{[k]} + P(\eta)_{[k]} + P(\theta)_{[k]} = 1$ <sup>2</sup>As expected, $P(A)_{[k]} + P(A)_{[k]} + P(\Gamma)_{[k]} + P(\Delta)_{[k]} + P(E)_{[k]} + P(Z)_{[k]} + P(H)_{[k]} + P(\Theta)_{[k]} = 1$

